# Graphical examples show why caution is required if using the coefficient of determination (R^2^) to interpret data for medical case reports

**DOI:** 10.1101/2025.08.18.670747

**Authors:** Thomas J. Hurr

## Abstract

A patient with a medical condition can have medical tests or symptoms scored that generate numerical results before a treatment, during a treatment or after a treatment, usually over several days, to determine if any benefits have occurred. The changes in the numerical measurements or scores over time can be readily plotted using computer software to show an equation for the line of best fit for either linear or log equations, together with the coefficient of determination (R^2^). Despite the ease of generating this type of graphical representations caution is required in interpreting the R^2^ value with reference to medical case reports.

To understand why this is so, at a basic level, four scenarios using hypothetical patient scores were used to generate scatter plots showing the equation for the line of best fit and R^2^ values with comparison to the average and standard deviation (SD) values. The graphical examples are used to supplement the more complex mathematical and statistical explanations and choice for effect measures that are available.

It was found R^2^ values for log equations for the line of best fit did not follow a trend with increasing treatment days. For linear equations, higher R^2^ value may not necessarily correspond to a lower standard deviation (SD) value for the averaged scores. The R^2^ value can be influenced by the day on which the scores were recorded, despite the equivalence of the average scores and SD values. R^2^ values may not indicate the strength of a treatment benefit or the magnitude of scatter between data sets. Score averaging can increase R^2^ values, while average values remain the same but with the SD value decreasing.

The graphical examples shown provide an explanation why line graphs may be the simplest and best option for reporting, particularly non-linear numerical data, in case reports.

**Graphical Abstract:** Graphical examples of the line of best fit and R^2^ values from hypothetical patient scores are compared with average (Av.) and standard deviation (SD) values
**A**. From the line of best fit, Patient 1 has a higher R^2^ value than Patient 2 even though the average score has a higher SD value. **B**. Patient 1 records scores on days 6 and 7 and Patient 2 records the same scores on days 9 and 10, yet Patient 1 has a higher R^2^ value for the line of best fit despite the scores average and SD values being the same. **C**. For Patients 1 and 2, the R^2^ values for the line of best fit are the same, despite the score averages and SD values being different and show R^2^ values do not predict a treatment benefit or allow a comparison of the magnitude of a benefit between data sets. **D**. Averaging daily scores removes scatter, increasing R^2^ values however the average scores remain the same, but the SD value (± 0.51) was reduced despite an identical slope and intercept for the line of best fit.

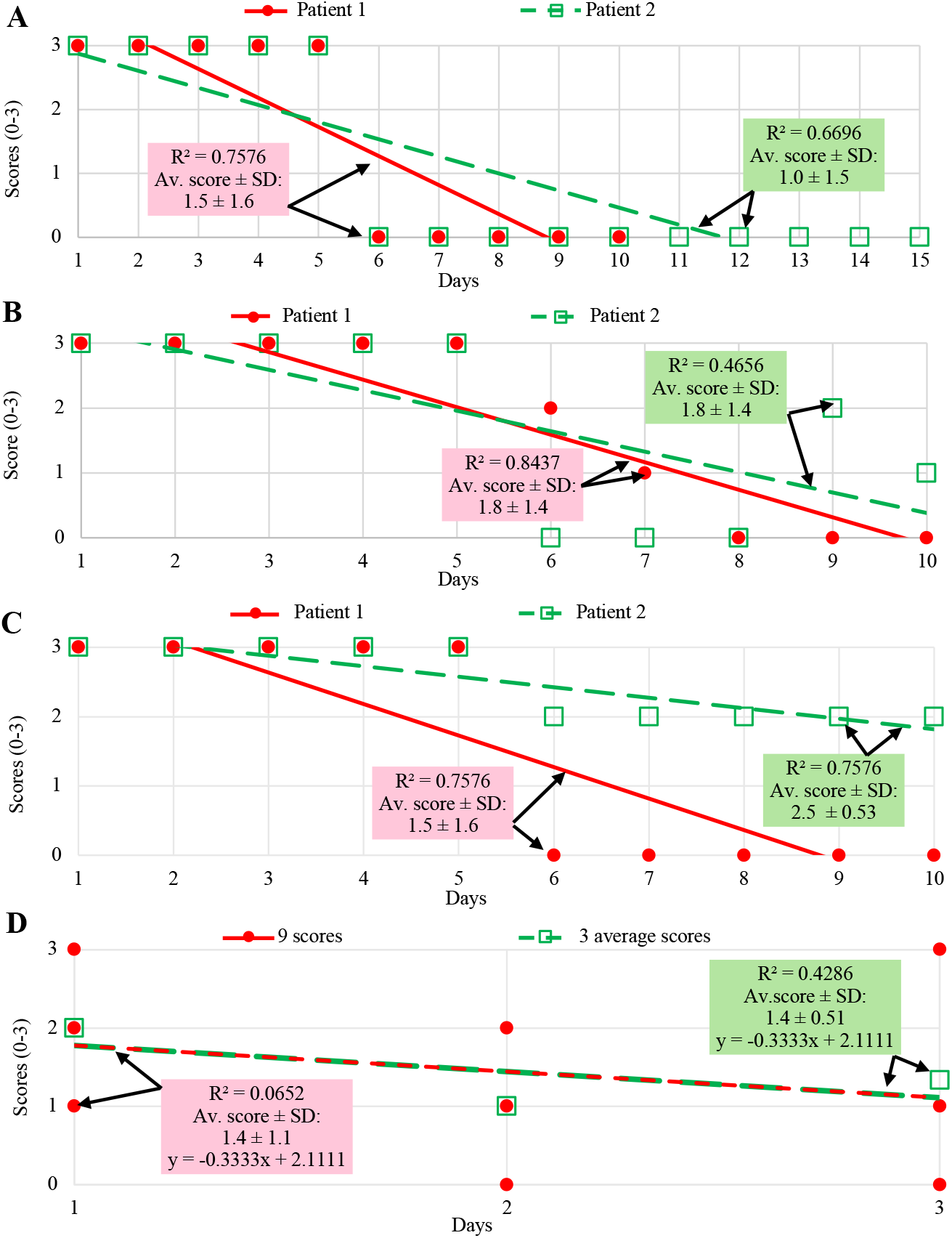

## 1.0 Introduction

Patients with a medical condition can have numerical measurements or symptoms scored before a treatment, during a treatment or after a treatment, to determine if any benefits have occurred. The benefits can depend on the treatment and so considered the dependant variable with benefits occurring over time, as the independent variable [1-3]. The results can generate a scatter plot and using readily available computer programs, the line of best fit for either linear or log equations and the coefficient of determination (R^2^) found [1-3]. The R^2^ value indicates how well the data fits (goodness of fit) the equation generated and has a value typically between 0-1, with a value of 1 indicating a perfect fit to a linear equation [1-3].

For linear equations, in the data set (x_i_,y_i_), the line of best fit takes the form y = mx + c, with m the gradient or slope of the line and c a constant such that when x = 0, the intercept for the y axis is (0,c). This equation can be used to predict a previously unknown value for y if a value for x is known, by substitution into the equation. It is now up to the observer to interpret whether this an appropriate way to measure the effect and what this means in relation to the data set under investigation.

Anscombe (1973) reported that graphs help us perceive, appreciate and understand data and are essential in statistical analysis [3]. He showed, using 4 data sets, 4 very different graphical representations of data with the same value of R^2^ = 0.667, a normal scatter plot, a scatter plot with a curve, a scatter plot with one point far from the line of best fit and a scatter plot with one point far from the other points and warned that sometimes one data point can play a critical role in the data analysis [3].

Schober et al. (2018) showed 3 scatter plots with a Pearson correlation coefficient of r = 0.84 all with different graphical appearances and one plot with a clear correlation between the points but with a correlation coefficient close to 0 (r = −0.05) [4]. When to use r or R^2^ was considered with r x r = R^2^ and it was suggested results should be shown graphically and inspected to check the correlations and not rely on numerical values of r or R^2^ alone [4].

In pharmacological and biochemical research, it has been reported that an evaluation of R^2^ was an inadequate measure for nonlinear models, however, were still being used frequently in the literature [5].

A vast literature is available with complex mathematical explanations for the meaning, calculation and use of R^2^ values including for use in clinical medicine, with guidelines available in choosing effect measures and computing estimates of effect [2,4-7].

The definition of key words used in this report include dispersion, as distribute or spread over a wide areas; scatter, as cover a surface with objects thrown or spread randomly over it and spread, as open out something so as to extend its surface area, width or length or disperse over an area, suggesting dispersion, scatter and spread can be used interchangeably [11].

This report uses 4 scenarios of hypothetical patient scores to highlight graphically, at a basic level, the difficulty and complexity of interpreting the equation for the line of best fit, R^2^ values and scatter in the data set under investigation [8-10]. A comparison to the mean and standard deviation (SD) values using scores that could arise, for example in medical case reports, are also given [8-10].

For the 4 scenarios, patients scores range from 0 - 3 with 0 being no symptoms, 1 being mild symptoms, 2 being moderate symptoms and 3 being significant symptoms.

## 2.0 Results and Discussion

### 2.1 Scenario 1: Linear or log equations for the line of best fit

In Scenario 1, 3 patients estimate the scores for their symptoms over each of the previous 5 days, (days 1-5) and all 3 patients have the same symptom score of 3, for each of the 5 days and then:

Patient 1 begins the treatment for 5 days, days 6-10 and records the scores of 0 over each of the 5 days,

Patient 2 begins the treatment for 10 days, days 6-15 and records the scores of 0 over each of the 10 days,

Patient 3 begins the treatment for 35 days, days 6-40 and records the scores of 0 over each of the 35 days.

The results are plotted using either linear or log equations to determine the line of best fit and R^2^ values, Fig. 1, Table 1. Appendix 1 shows how to estimate a line of best fit and calculate the R^2^ value for the data set shown in Fig. 1A.

**Table 1.** Results for Scenario 1.

| | Linear equation<br>$y = m x + c$ | | log equation<br>$y = \ln x + c$ | Average score and standard deviation (SD) | |
| --- | --- | --- | --- | --- | --- |
| | Slope m | $R^2$ | $R^2$ | Over all days | Over treatment days only |
| Patient 1 | -0.4545 | 0.7576 | 0.6322 | $1.5 \pm 1.6$ (days 0-10) | $0 \pm 0$ (days 6-10) |
| Patient 2 | -0.2679 | 0.6696 | 0.7135 | $1.0 \pm 1.5$ (days 0-15) | $0 \pm 0$ (days 6-15) |
| Patient 3 | -0.0492 | 0.3283 | 0.6227 | $0.38 \pm 1.0$ (days 0-40) | $0 \pm 0$ (days 6-40) |

**Figure 1.**
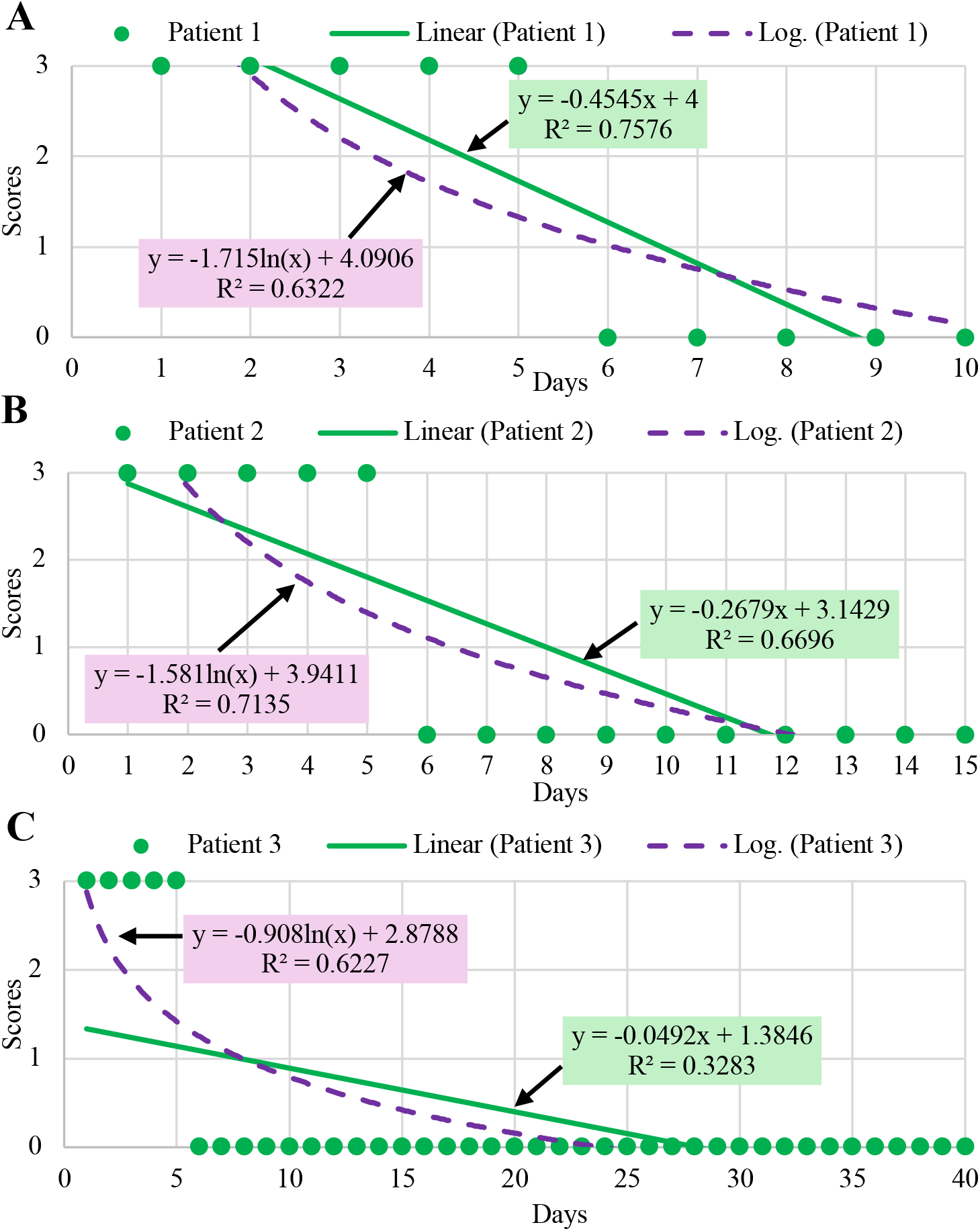
**A.** Patient 1 has 5 days without treatment followed by 5 days with treatment, over a total of 10 days **B**. Patient 2 has 5 days without treatment followed by 10 days with treatment, over a total of 15 days. **C**. Patient 3 has 5 days without treatment followed by 35 days with treatment, over a total of 40 days.

The log equations for the line of best fit for the 3 patients show a consistent change in the shape of the curved line of best fit with the increasing number of days with a score of 0, but the R^2^ values for 10 days were R^2^ = 0.6322, for 15 days R^2^ = 0.7135 for 40 days R^2^ = 0.6227 for 40 days without showing an expected trend, resulting in inconclusive results and so should possibly be avoided, Fig. 1, Table 1, [5].

For the linear equations it is now up to the observer to compare the results for the 3 patients and interpret the scenario, Figs. 1-2, Table 1. It is thought this scenario shows:

- from Fig. 1, for the increasing days where the score is 0, the decreasing R^2^ values indicate a decreasing goodness of fit to the linear equations and suggest that non-linear equations may better describe the data,
- from Table 1, the decreasing R^2^ value indicates increasing dispersion or scatter in the data relative to the line of best fit,
- from Table 1, Fig. 2, for the increasing days where the score is 0, the SD decreased indicate a decreasing dispersion in the data relative to the mean values.

**Figure 2.**
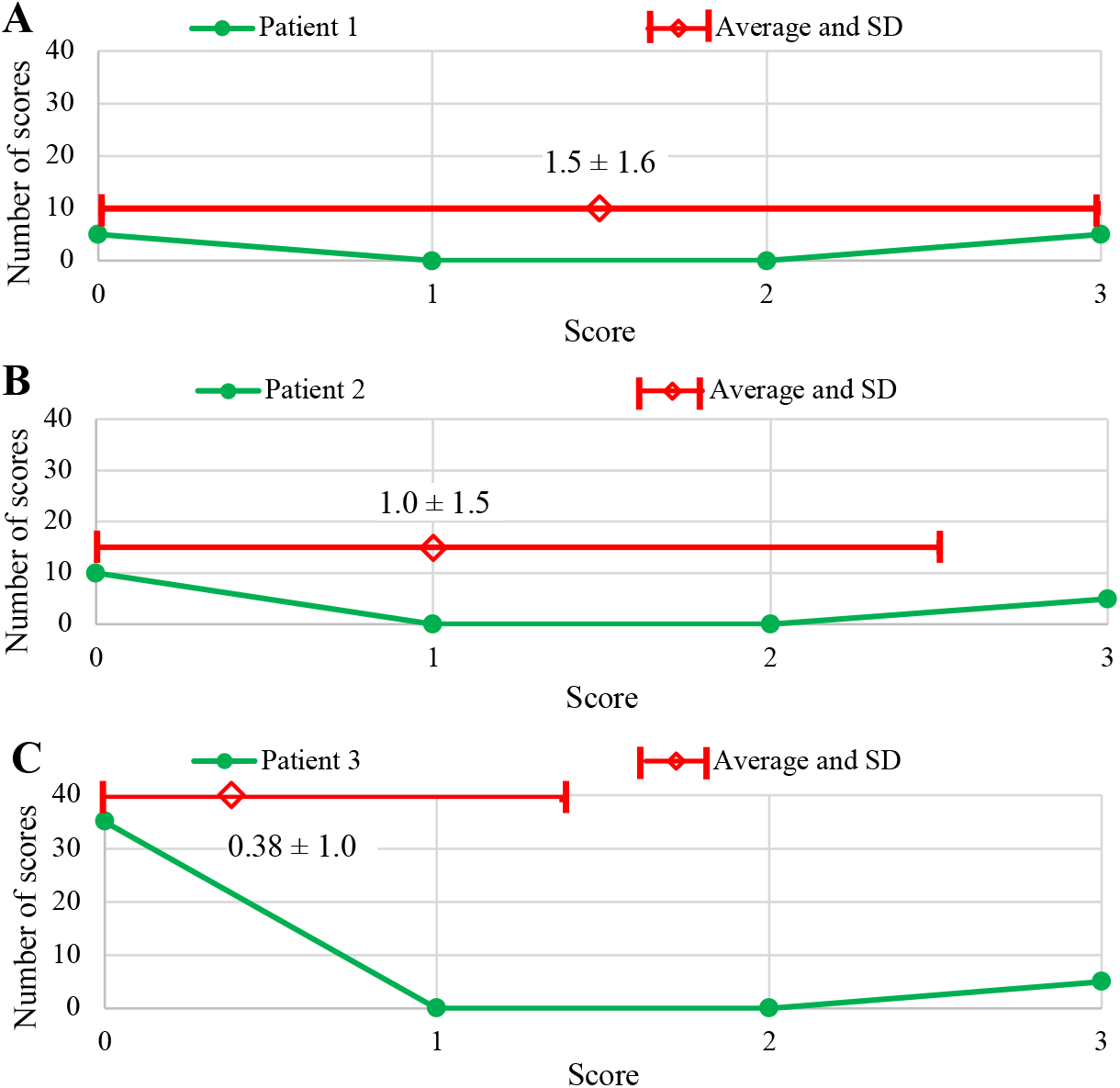
**A,** Scores (0-3) vs the number of scores (score frequency) recorded over the 10 days with average and SD values for Patient 1. **B**, Scores (0-3) vs the number of scores (score frequency) recorded over the 15 days with average and SD values for Patient 2. **C** Scores (0-3) vs the number of scores (score frequency) recorded over the 40 days with average and SD values for Patient 3. The average and SD values for Patients 1-3 over the scoring range (0-3) show that as the number of days with a score of 0 increase, the average score and SD values reduce, showing a decrease in the dispersion of the data set relative to the average values.

The R^2^ value is calculated from a ratio of the y values such that any dimensions the y values may have had will cancel and so become dimensionless (Appendix 1). The SD values are in the same units as the data itself, for example velocity (v) in metres (m) per second (s) could be reported as v = 5±1 m/s with 2 dimensions as length (L) and time (T). Scores used in this report are simply a number without dimension but scores per day (score/day) have the dimensions of scores/time which can be written as 1/T, with a dimension of 1. The R^2^ and average with SD values described above, although they are both used to describe the scatter in the data, have different dimensions in this example. The example above shows a trend of decreasing R^2^ values with increasing days with a score of 0, indicating increased scatter, while the mean and SD values are reducing, indicating decreasing scatter in the data.

In summary, for the linear equations shown above, R^2^ values indicate increasing dispersion in the data from Patient 1 to Patient 3 (R^2^: 0.7576−0.3283), relative to the line of best fit, while in contrast the SD values for the means indicate a decreasing dispersion in the data (SD:1.6-1.0).

### 2.2 Scenario 2: The timing of occurrence for scores influences both the slope and R^2^ values

In the Scenario 2, Patients 1 and 2 estimate the scores for their symptoms over each of the previous 5 days (days 1-5) with both having the same symptom scores of 3 for each of the first 5 days and then:

Patient 1 begins the treatment for 5 days, days 6-10 and has symptoms for the first 2 days, day 6 (score of 2) and day 7 (score of 1) with no symptoms for days 8-10,

Patient 2 also begins the treatment for 5 days, days 6-10 and has the same symptom scores but for the last 2 days, day 9 (score of 2) and 10 (score of 1) with no symptoms for days 6-8.

The results are shown using only linear equations, as the use of log equations in Scenario 1 gave R^2^ values that were inconclusive, Fig. 3. The average scores and SD values are also shown in Table 2, Fig. 4.

**Table 2.** The results for Scenario 2.

| | Linear equation<br>$y = m x + c$ over all<br>days 1-10 (10 days) | | Linear equation<br>$y = m x + c$ over treatment<br>days only days 6-10 (5 days) | | Average score and standard<br>deviation (SD) | |
| --- | --- | --- | --- | --- | --- | --- |
| | Slope m | $R^2$ | Slope m | $R^2$ | Over all<br>days (1-10) | Over treatment<br>days only (6-10) |
| Patient 1 | -0.4242 | 0.8437 | -0.5 | 0.7813 | $1.8 \pm 1.4$ | $0.60 \pm 0.89$ |
| Patient 2 | -0.3152 | 0.4656 | 0.4 | 0.5 | $1.8 \pm 1.4$ | $0.60 \pm 0.89$ |

**Figure 3.**
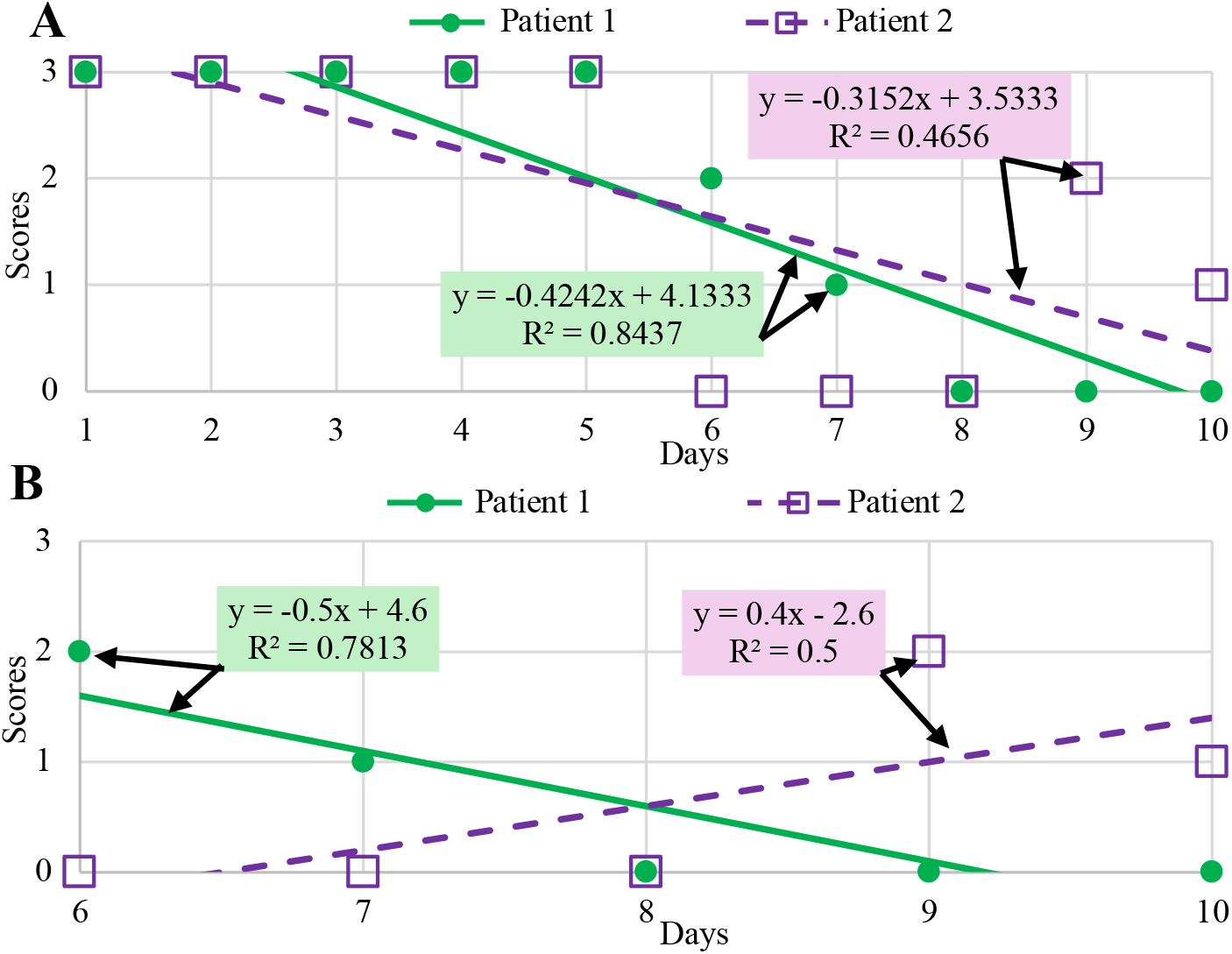
**A**. Results for Patient 1 and 2 over days 1-5 without treatment followed by days 6-10 (5 days) with treatment, both with the same scores, but on different days. **B**. Results for Patients 1 and 2 over days 6-10, the treatment days only.

**Figure 4.**
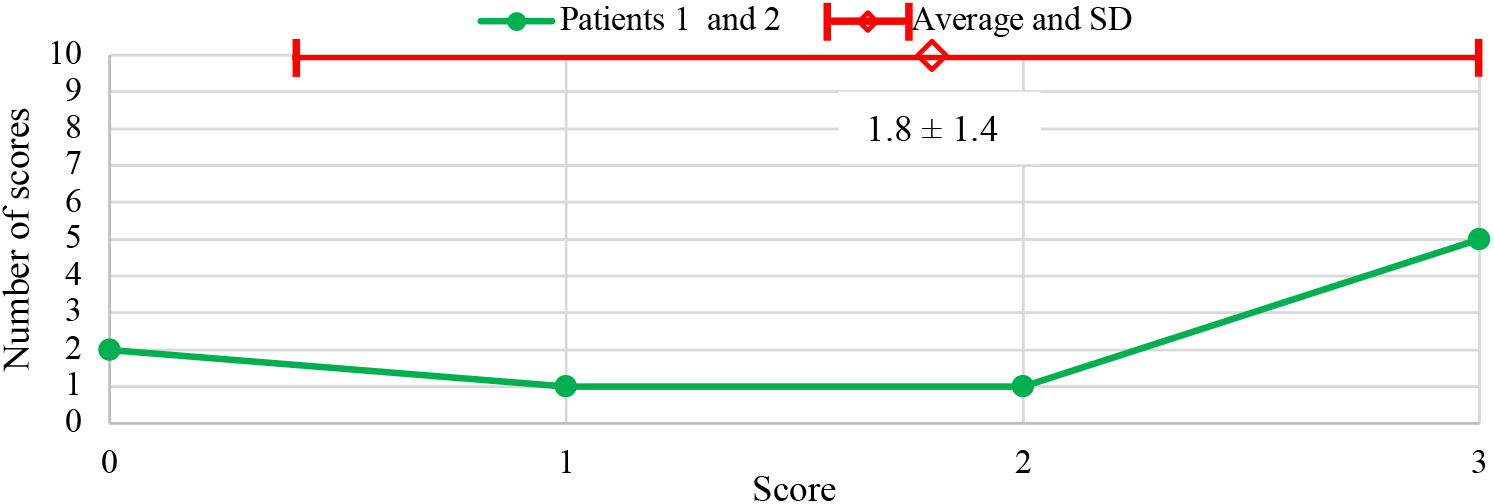
Patient scores (0-3) vs the number of scores (score frequency) recorded over the 10 days with the average and SD values. Patients 1 and 2 have the same average and SD values but the graph shows no information of how the scores changed over the 10 days as shown in Fig. 3.

It is now up to the observer to compare the results for the 2 patients and interpret the Scenario, Figs. 3,4 and Table 2. It is thought this Scenario shows:

- the slope of the equation for the line for Patient 1 is more negative, with a value of - 0.4242 compared to Patient 2, with a slope of −0.3152, which could be thought to suggest symptoms for Patient 1 decline more rapidly than Patient 2, but symptoms for Patient 2 decline more rapidly, being 0 on day 6-8 before increasing,
- a comparison of the R^2^ values could be thought to suggest the benefits were greater for Patient 1, with a better “goodness of fit” even though both patients had the same symptom scores (1 and 2), over the 5 treatment days, just on different days with the same average and SD score values,
- the R^2^ values indicate the dispersion or scatter of the scores relative to the line of best fit, with a better fit to the scores for Patient 1 (R^2^ = 0.8437) than Patient 2 (R^2^ = 0.4656) but the average and SD values of 1.8 ± 1.4 for both patients show the dispersion or spread from the SD values was the same, Table 2, Fig. 4.

For the Scenario where the scores are plotted for the treatment days only (days 6-10), as shown in Fig 3B, Table 2, it is thought this scenario shows:

- from a comparison of the R^2^ values, scores for Patient 1 are a better fit to the linear equation than Patient 2,
- the line of best fit has a positive slope for Patient 2, showing a trend towards a loss of treatment benefit over time from the initial score of 0, over the 5 days, compared to Patient 1 where results show a negative slope for the line of best fit with an initially delayed benefit, but showing a score of 0 towards the end of the 5 days of treatment, Fig. 3B,
- the slopes (one positive and one negative) and R^2^ values relative to the line of best fit for Patient 1 and 2, are not the same, despite both patients showing the same average score and SD values of 0.60 ± 0.89, Table 2.

In summary, for the same number and value of symptom scores over 10 days, or over only the 5 days treatment, the timing of symptom as on days 6, 7 or on days 9, 10 can change the slope of the line of best fit and the R^2^ values, even though the scores have the same average and SD values, Table 2, Figs. 3,4.

### 2.3 Scenario 3: R^2^ values do not indicate the strength of a treatment benefit or the magnitude of scatter between data sets

In Scenario 3, 2 patient estimates the scores for their symptoms over the previous 5 days, (days 1-5) and both have the same symptom scores of 3 for each of the 5 days then:

Patient 1 begins the treatment for 5 days, days 6-10, and has no symptoms with a score of 0 is recorded for each day.

Patient 2 also begins the treatment for 5 days, days 6-10 but has symptoms each day, with a score of 2 recorded.

It is now up to the observer to compare the results for the 2 patients and interpret the Scenario, Figs. 5,6. It is thought this scenario shows:

**Figure 5.**
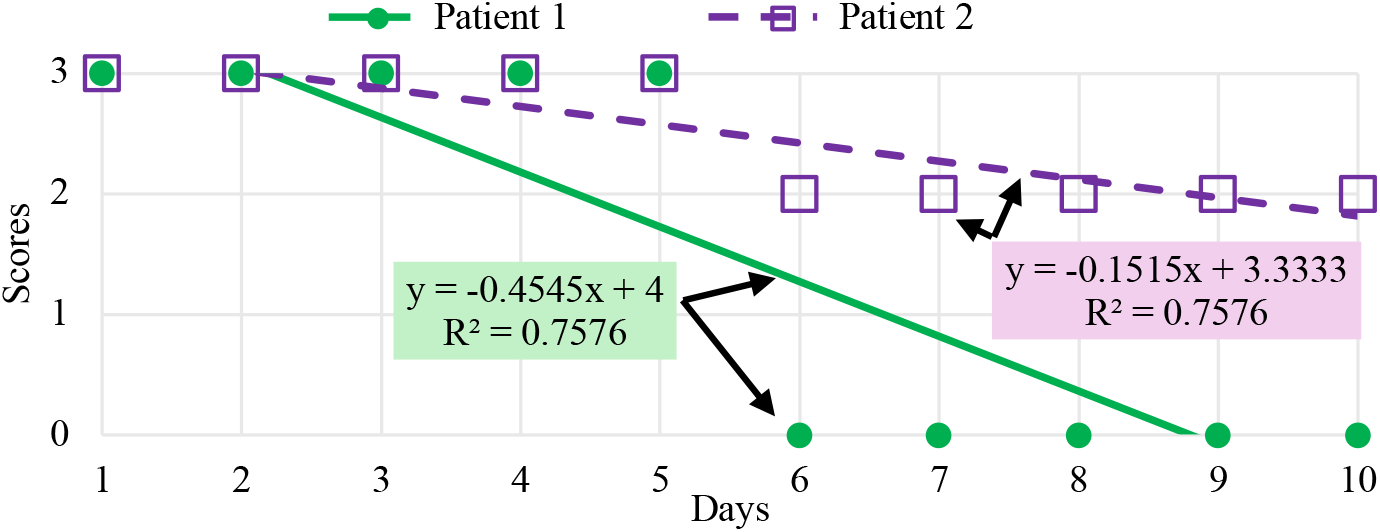
Scenario 3. Results from Patient 1 and 2 over 10 days, with estimated scores 5 days prior to treatment days 1-5 and then scores after treatment, days 6-10.

**Figure 6.**
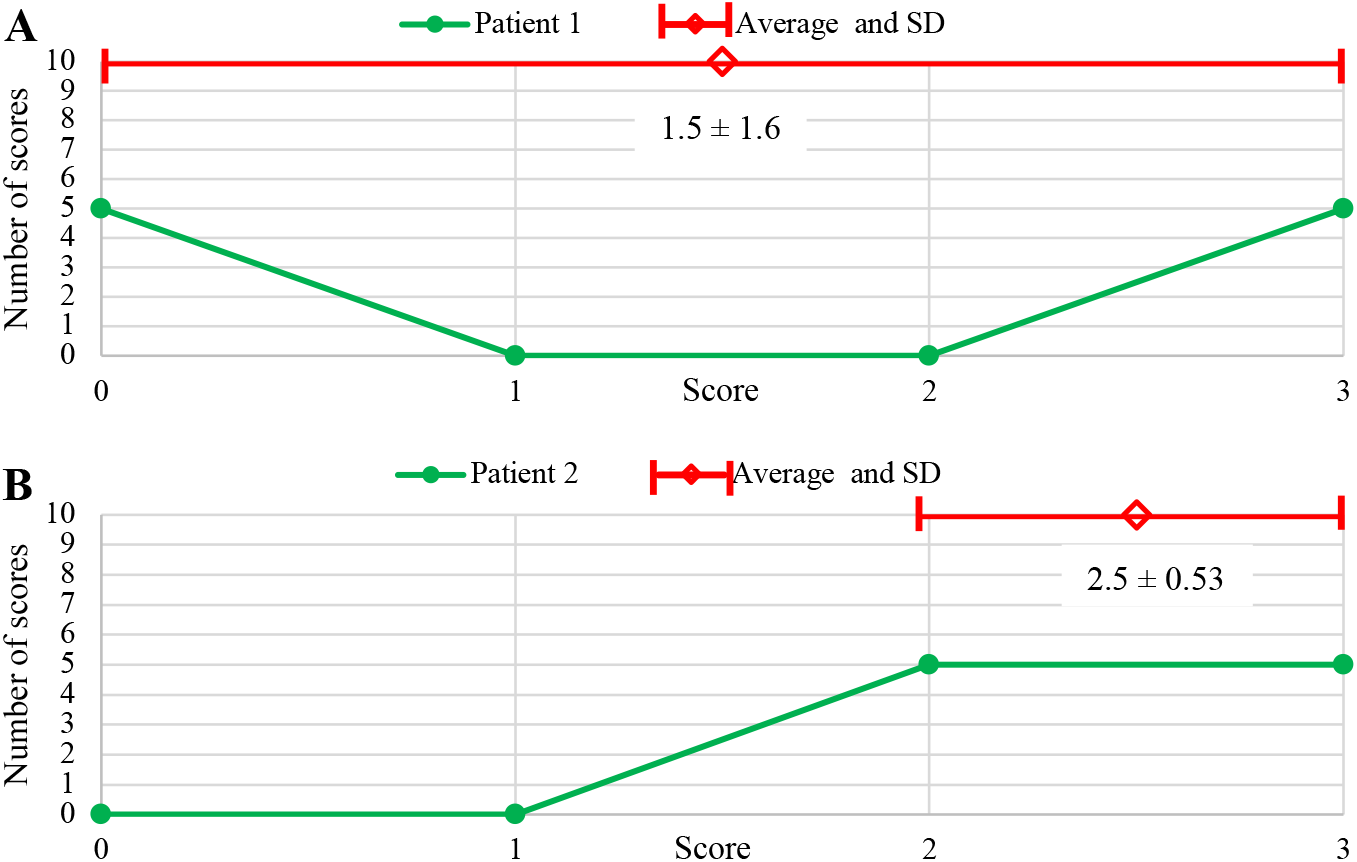
Patient scores (0-3) vs the number of scores (score frequency) recorded over the 10 days. Patients 1 and 2 have the same R^2^ values but different average and SD values.

- the slope of both the linear equations decreases with treatment time, suggesting a treatment benefit for both patients, with a greater benefit for Patient 1, as shown by a decrease in the average scores, Fig. 5, Table 3,
- the R^2^ values are the same for both Patients 1 and 2 showing R^2^ values indicate the “goodness of fit” but not strength of a treatment benefit as Patient 1 has a greater treatment benefit with a lower average score than Patient 2 over the 10 days,
- it could be suggested that Patient 1 has more dispersion in the data than Patient 2 as the data points are more dispersed or spread out, being between scores of 3 and 0 rather than between scores of 3 and 2, but the R^2^ value does not allow a comparison of the magnitude of scatter between data sets, only the relative degree of scatter to the line of best fit, as shown in Fig. 5 (Appendix 2),
- the R^2^ values can be the same despite a different in the slopes for the equation for the line of best fit,
- Patient 1 has a greater treatment benefit than Patient 2, with a lower average score but with a larger SD value, but this is not reflected by the R^2^ values (both R^2^ = 0.7576), Fig. 6, Table 3.

In summary, R^2^ represents the relative dispersion within a data set in relation to the line of best fit but not necessarily the magnitude of the dispersion itself, to allow a comparison to be made between different data sets.

**Table 3.**
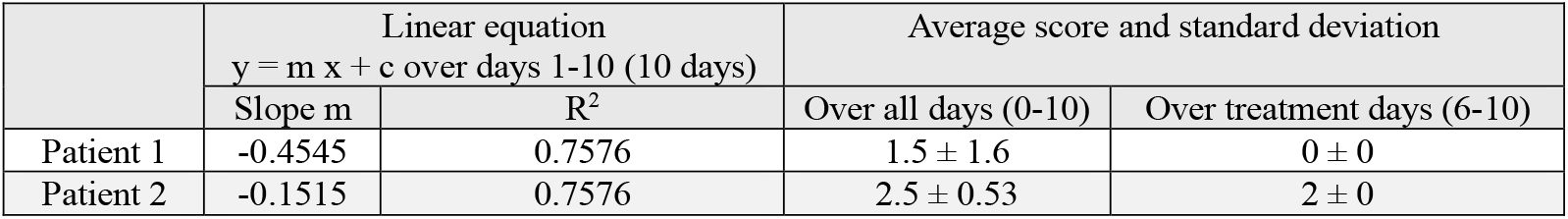
The results for Scenario 3 from Fig. 5.

| | Linear equation<br>$y = m x + c$ over days 1-10 (10 days) | | Average score and standard deviation | |
| --- | --- | --- | --- | --- |
| | Slope m | $R^2$ | Over all days (0-10) | Over treatment days (6-10) |
| Patient 1 | -0.4545 | 0.7576 | $1.5 \pm 1.6$ | $0 \pm 0$ |
| Patient 2 | -0.1515 | 0.7576 | $2.5 \pm 0.53$ | $2 \pm 0$ |

### 2.4 Scenario 4: Averaging scores removes scatter and increases R^2^ values

In Scenario 4, a patient estimates the scores (0-3) for symptoms three times a day over 3 days, to determine a baseline, before a treatment could begin, Table 4. The scores are either plotted as the 9 separate scores over the 3 days or as 3 daily averaged scores, Fig. 7.

**Table 4.** The patient scores symptoms 3 x per day, resulting in 9 scores. These scores can be averaged, to give 3 daily average scores, with average, daily average and SD scores given.

| Day | Scores 3x per day | Average of the 9 scores over 3 days and SD | Average of the 3 daily scores | Average for the 3 averaged daily scores and SD |
| --- | --- | --- | --- | --- |
| 1 | 1<br>2<br>3 | (1+2+3+0+2+1+0+1+3)/9<br>= 13/9<br>= 1.4 ± 1.1 | (1+2+3)/3<br>= 2 | (2+1+1.333)/3<br>= 4.333/3<br>= 1.4 ± 0.51 |
| 2 | 0<br>2<br>1 |  | (0+2+1)/3<br>= 1 |  |
| 3 | 0<br>1<br>3 |  | (0+1+3)/3<br>= 1.333 |  |

**Figure 7.**
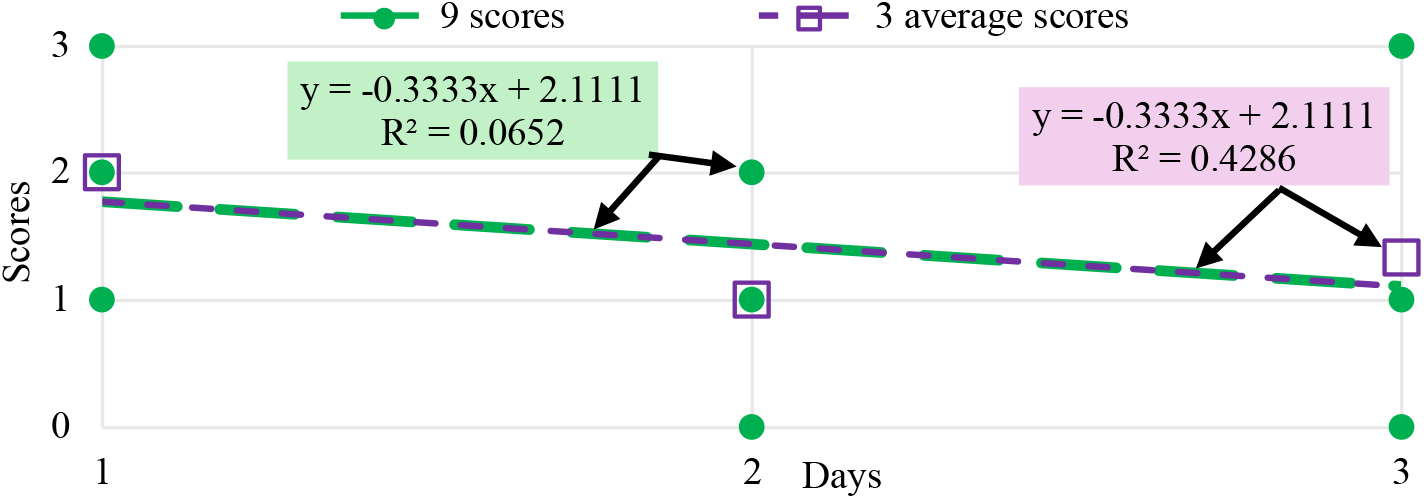
Results from the patient scores over days 1-3, with the line of best fit and R^2^ value for all 9 scores over the 3 days or with scores averaged daily to give 3 scores over the 3 days, from Table 4.

It is now up to the observer to compare the results for the patient and interpret the scenario, Fig. 7, Tables 4,5. It is thought this scenario shows:

**Table 5.** The patient scores symptoms 3 x per day, resulting in 9 scores for days 1-3. The 3 daily scores over the 3 days (1-3) are also averaged for each day to give 3 averaged daily scores.

| | Linear equation $y = m x + c$ with all 9 scores for days 1-3 | | | Linear equation $y = m x + c$ with daily scores averaged resulting in 3 scores for days 1-3 | | |
| --- | --- | --- | --- | --- | --- | --- |
| | Slope m | Intercept c | $R^2$ | Slope m | Intercept c | $R^2$ |
| Patient 1 | -0.3333 | 2.1111 | 0.0652 | -0.3333 | 2.1111 | 0.4286 |

- some of the scatter is removed from the data when average scores are used, resulting in a significantly higher value for R^2^ (0.4286 rather than 0.0652),
- plotting all 9 scores or only the 3 averaged scores result in the same average score of 1.4, but the SD for the averaged score is lower at 0.51, as scatter was removed, Table 4,
- averaging scores does not change the slope and the intercept for the line of best fit, Fig. 7.

In summary, averaging scores and using these scores in a scatter plot, will result in reduced scatter, the same equation for the line of best fit and an increased R^2^ value, possibly leading to a conclusion that a treatment has a greater benefit than may otherwise be predicted.

### 2.5 A simple line graph may be the best option if the data set is non-linear

A simple line connecting all the data points is shown for Scenario 2, Patient 1 and 2, without the lines of best fit or R^2^ values with a loss of the predictive power of equations to describe the data, but the trends are still apparent, Fig. 8.

**Figure 8.**
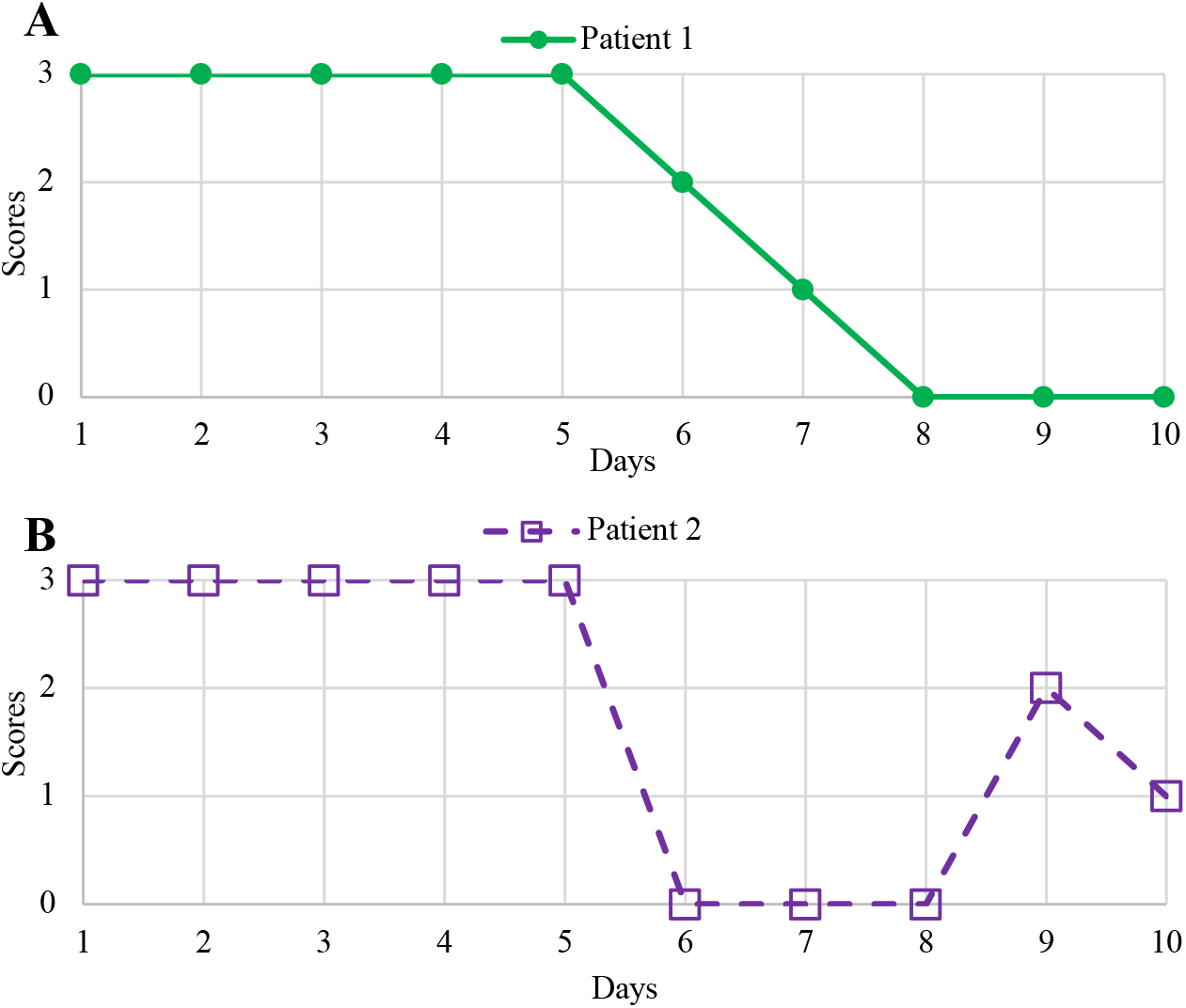
**A, B.** A simple connecting line of symptom scores for Patients 1 and 2 in Scenario 2 from Fig. 2A, replacing the line of best fit.

It is now up to the observer to compare the results for the 2 patients and interpret the scenario, Fig. 8. It is thought this Scenario shows:

- Patient 1 shows a slower response to treatment and Patient 2, a fast initial response, but this was not sustained for days 9 and 10.
- Both patients show the same improvement in symptoms from an average pre-treatment score of 3 to 1.8 ± 1.4 over the 10 days or from a pre-treatment score of 3 to a score of 0.6 ± 0.89 over the 5 days of treatment, Table 2.

In summary, a simple line graph may be the best option, together with the average and SD values, especially for medical case reports where small data sets may only be available and control over the variables that may arise in the case, may not be possible.

## 3.0 Conclusion

Dispersion, as scatter in the data set with variables (x_i_,y_i_), can be indicated by the determination of the R^2^ value for a linear equation of the form y = mx + c. It is now up to the observer to interpret whether this an appropriate way to measure the effects and what this means in relation to the data set under investigation.

It was shown that the R^2^ value for the line of best fit may not show the same trend to describe the scatter in the data set, as the average and SD value.

It was also shown, if scatter is defined as the magnitude of the separation between data points on a scatter plot, the R^2^ value only represents the relative scatter to the line of best fit for one data set and not the magnitude of scatter between data sets.

A line connecting the data points may be the simplest and best option if the data set is not linear, unless caution is used in the interpretation of the R^2^ values, with the average scores and SD values given.

## Appendix 1

### Estimating the line of best fit and calculating the R^2^ value for Figure 1A

#### 1.1. Estimating the line of best fit through the data points

Step 1: Estimate the line of best fit by drawing a line through the data points (scores) with the line showing some scores on both sides and intersecting at least 2 scores, for the data from Fig. 1A, shown in Fig. 9A.

**Figure 9.**
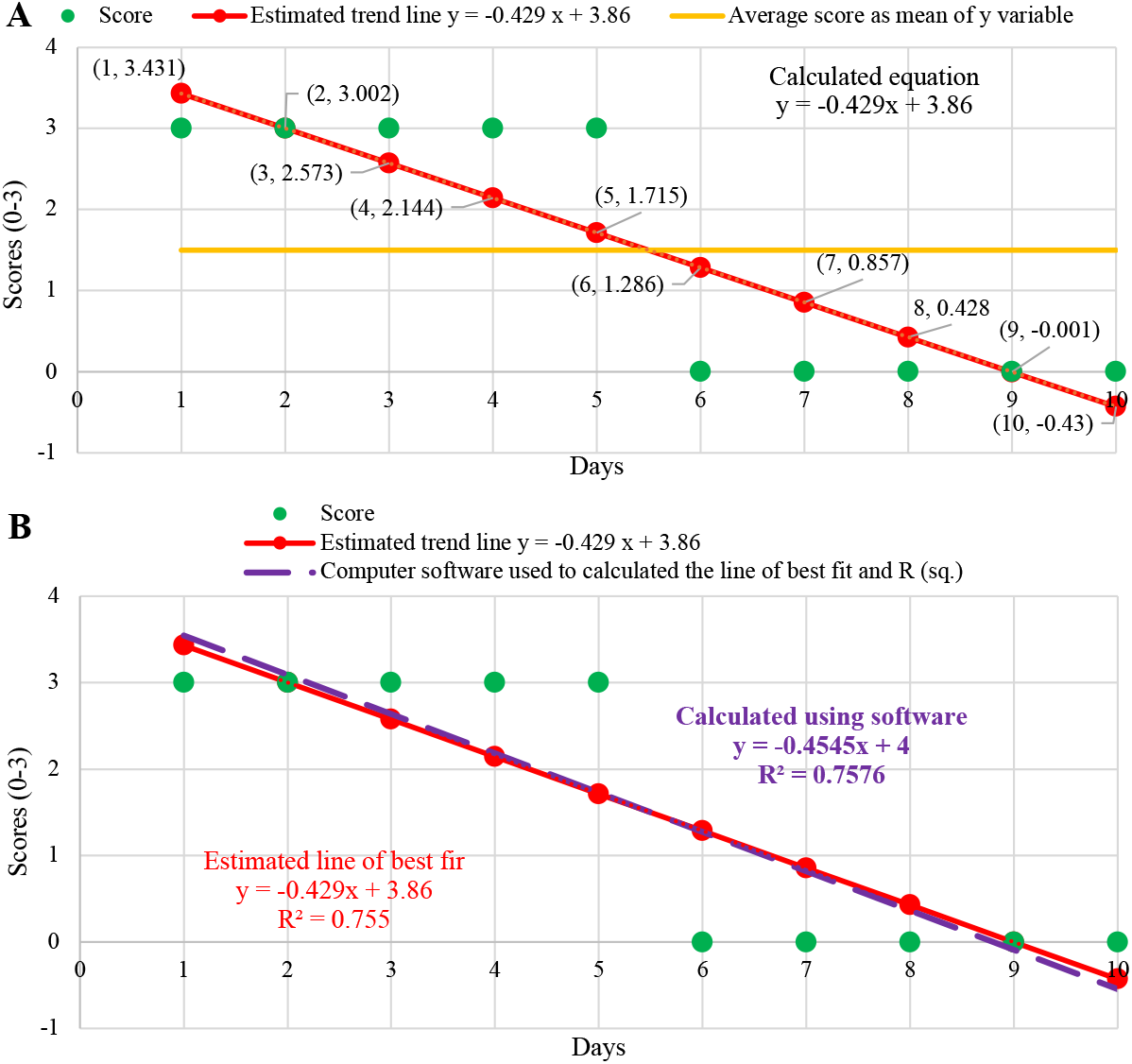
**A.** To estimate the equation for line of best fit, drawn a line between the scores shown in Fig. 1A and calculate the equation for the line, for example y = −0.429x + 3.68 then put the x values into the equation to determine the y values for the estimated line. Then use eq. 2 to calculate the value of R^2^ = 0.755 as shown in Table 6. **B**. The equation for the line of best fit calculated from computer software as y = −0.4545x + 4 with R^2^ = 0.7576 is similar to the estimated equation and calculated R^2^ = 0.755 above.

Step 2: Choose 2 data points on the estimated line for example (2,3) as (x_1_,y_1_) and (9,0) as (x_2_,y_2_) and calculate the slope for the line as:

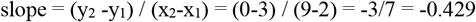

and now the estimated equation for the line is y = −0.429x + c.

**Table 6.** The estimated line of best fit is used to calculate R^2^ from eqs. 1,2 for the scores shown in Fig. 1A.

| Days<br>(x axis) | Scores<br>(y axis<br>scores as<br>$y_i$) | Calculated scores<br>from the estimated<br>equation of the line of<br>best fit $y = -0.429x + 3.86$ using x values to<br>calculated ($\hat{y}$) | Mean of<br>the y or<br>$y_i$ values<br>from the<br>scores<br>($\bar{y}$) | Scores ($y_i$) subtract<br>calculated scores ($\hat{y}$)<br>from the estimated<br>equation squared<br>($y_i - \hat{y}$)<sup>2</sup> | Scores ($y_i$)<br>subtract mean<br>of scores ($\bar{y}$)<br>squared<br>($y_i - \bar{y}$)<sup>2</sup> | (SS<sub>res</sub>)/<br>SS<sub>total</sub> | $R^2 = 1 -$<br>(SS<sub>res</sub>)/SS<sub>total</sub> |
| --- | --- | --- | --- | --- | --- | --- | --- |
| 1 | 3 | $\hat{y} = -0.429 \times 1 + 3.86$<br>$= 3.431^a$ | 1.5 | $(3 - 3.431)^2 = 0.186^a$ | $(3 - 1.5)^2 = 2.25^a$ | | |
| 2 | 3 | $\hat{y} = -0.429 \times 2 + 3.86$<br>$= 3.002^a$ | 1.5 | $(3 - 3.002)^2 = 0.000004^a$ | $(3 - 1.5)^2 = 2.25^a$ | | |
| 3 | 3 | $\hat{y} = -0.429 \times 3 + 3.86$<br>$= 2.573^a$ | 1.5 | $(3 - 2.573)^2 = 0.182^a$ | 2.25 | | |
| 4 | 3 | 2.144 | 1.5 | 0.732 | 2.25 |  |  |
| 5 | 3 | 1.715 | 1.5 | 1.651 | 2.25 |  |  |
| 6 | 0 | 1.286 | 1.5 | 1.653 | $(0 - 1.5)^2 = 2.25^a$ | | |
| 7 | 0 | 0.857 | 1.5 | 0.734 | $(0 - 1.5)^2 = 2.25^a$ | | |
| 8 | 0 | 0.428 | 1.5 | 0.183 | 2.25 |  |  |
| 9 | 0 | -0.001 | 1.5 | $\approx 0$ | 2.25 | | |
| 10 | 0 | -0.43 | 1.5 | 1.85 | 2.25 |  |  |
| | Mean ($\bar{y}$)<br>$= 15/10 = 1.5$ | | | (SS<sub>res</sub>) = $\sum_{i=1}^n (y_i - \hat{y})^2 = 5.508$ | SS<sub>total</sub> = $\sum_{i=1}^n (y_i - \bar{y})^2 = 22.5$ | 5.508/22.5 = 0.2448 | 1 - 0.2448 = 0.755 |

Step 3: To calculate the intercept, choose a data point for example (9,0) and put this into the estimated equation, y = 0.429x + c to give:

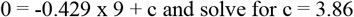

Step 4: Write the full equation for the estimated line of best fit:

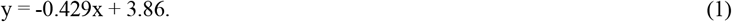

#### 1.2 Calculating R^2^ for the estimated line of best fit

The coefficient of determination R^2^ is calculated as:

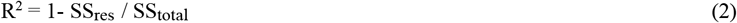

where 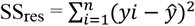 is the residual sum (*yi* − *ŷ*)^2^ of the squares as the sum of the squared difference between observed y values (scores) as *yi* in eq. 1 subtracted from ŷ, calculated using the predicted y values calculated from the estimated line of best fit (y = −0.429x + 3.86) using the x values (days 1,2,…10), 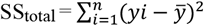 is the sum of the squared differences between observed variables *yi* (*scores*) subtracted from the mean of the observed values 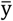, (the mean of the y values for all the scores), with n is the number of observations.

How to calculate the SS_res_ and SS_total_ values from eqs. 1,2, with the final calculated value of R^2^ = 0.755, is shown in Table 6, and found similar to that calculated by computer software of R^2^ = 0.7576.

## Appendix 2

### The R^2^ value indicates the relative scatter in the data set but does not allow a comparison of the magnitude of scatter between data sets

**Figure 10.**
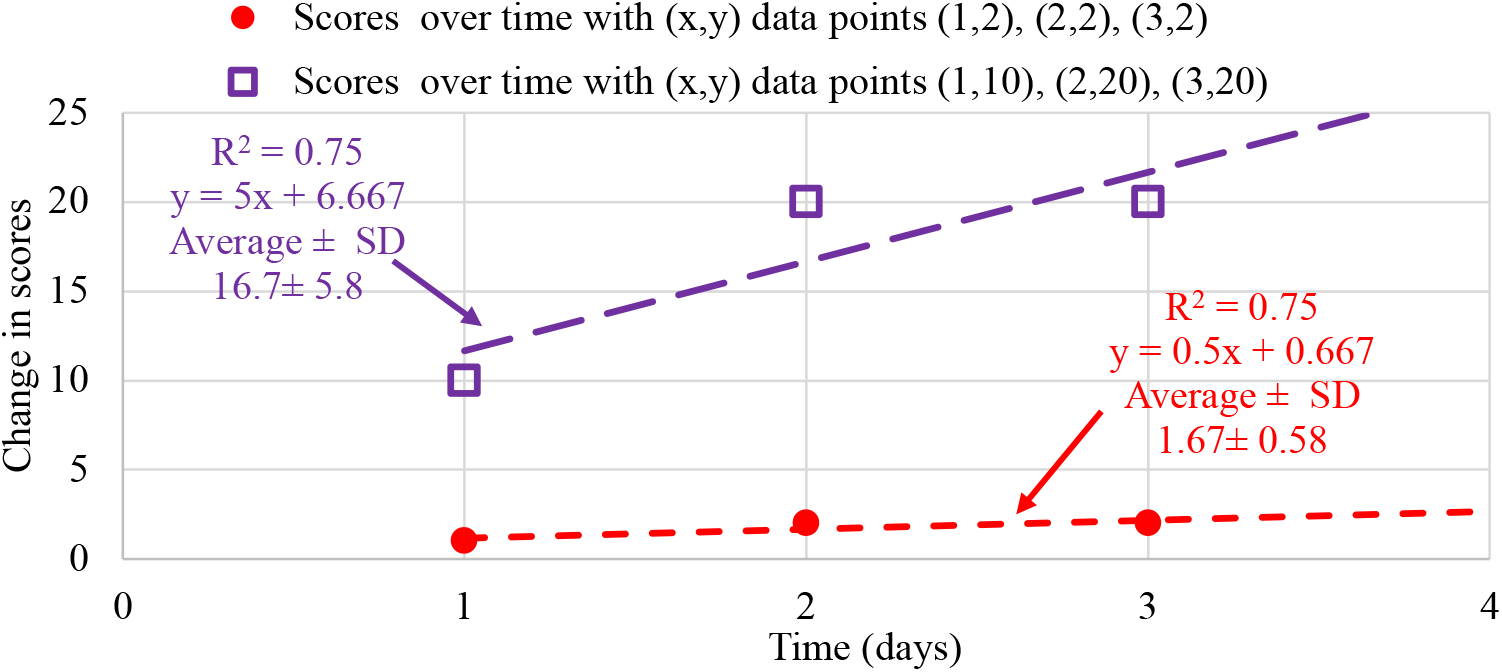
It could be thought that data points separated by 10 or 20 points may be more scattered apart from each other and have a greater magnitude of scatter than if separated by 1 or 2 points, as shown in the graph, but the R^2^ values are identical, showing R^2^ values only indicate the relative scatter from the line of best fit for the data set and not the magnitude of the scatter between data sets as does the average and SD values.

